# Discovery and *in vivo* characterization of novel TOG domain-containing proteins using *C. elegans*

**DOI:** 10.64898/2026.02.23.707544

**Authors:** Linnea C Wethekam, Taara Mittal, Amy Shaub Maddox

## Abstract

The proper assembly, architecture, and maintenance of microtubule, actin and other cytoskeletal networks require regulation by various polymer binding proteins. Microtubules, rely on both the tubulin building blocks, but also many tubulin- and microtubule binding proteins. TOG domain-containing proteins comprise one family of tubulin-binding proteins that regulate microtubule dynamics. Here we identify two previously uncharacterized TOG domain-containing proteins (TOD-1 and TOD-2) in the nematode, *C. elegans*. These proteins are unique in that they are members of the XMAP215 family but contain reduced numbers of TOG domains and, in one case, a divergent TOG domain. TOD-1 and TOD-2 are expressed in and contribute to the normal function of sperm. The single TOG domain of TOD-1 and both TOG domains of TOD-2 are predicted to bind free tubulin dimers and not microtubule lattice. Deletion of either *tod* gene resulted in an increased laying of unfertilized oocytes. Inspection of mutant hermaphrodites revealed a premature onset of sperm migration failure. Together, these findings suggest that *C. elegans* requires regulation of tubulin dimers and/or microtubules for sperm localization and function. The amoeboid movement of *C. elegans* sperm has been considered microtubule-independent, our results open a new avenue of research into their unique motility.

## Introduction

Cytoskeletal networks are built and remodeled to define the shape and function of each cell. Of the four families of cytoskeletal filaments, the microtubule cytoskeleton is fundamental to building cilia, mitotic spindles, the backbone of axons, and the tracks for dynein or kinesin driven transport. While α/β-tubulin heterodimers can assemble into microtubules spontaneously *in vitro*, within cells, the assembly and dynamics of microtubules are spatiotemporally regulated (Hyman et al., 1992; Mitchison & Kirschner, 1984). A large cohort of proteins binds microtubules to spatially modulate dynamic instability, including accelerating growth rate, preventing catastrophe, or promoting rescue events (Al-Bassam et al., 2010; Brouhard et al., 2008; Farmer et al., 2021; Lawrence et al., 2018; Majumdar et al., 2018; Slep & Vale, 2007; Tymanskyj & Ma, 2019). These proteins are called microtubule associated proteins with one subgroup called microtubule plus-end binding proteins. Microtubule plus-end binding proteins canonically bind to and modulate microtubule plus end dynamics, changing growth rate, shrinkage rate or the frequency with which microtubules switch between growth and shrinkage.

One class of plus-end binding proteins is the TOG (tumor overexpressed gene) domain-containing superfamily. There are three major families of TOG domain-containing proteins: the XMAP215 family, the CLASP family, and the Crescerin/TOGARAM family, each with different roles in regulating microtubule networks (Akhmanova et al., 2001; Cassimeris & Morabito, 2004; Das et al., 2015; Leano et al., 2013). XMAP215 and homologues help nucleate microtubule growth from the γ-tubulin ring complex and serve as polymerases, accelerating growth at the plus end (Ayaz et al., 2012; Brouhard et al., 2008; Gunzelmann et al., 2018; Widlund et al., 2011). CLASP family proteins are rescue factors (Al-Bassam et al., 2010; Gareil et al., 2023; Lawrence et al., 2018; Majumdar et al., 2018; Rodgers et al., 2023). Crescerin family proteins are responsible for the slow and sustained growth of microtubules in cilia (Das et al., 2015; Saunders et al., 2025). TOG domains are α-solenoid structures composed of multiple HEAT repeats (Al-Bassam et al., 2007; Byrnes & Slep, 2017; Leano & Slep, 2019; Slep & Vale, 2007). Loops between the HEAT repeat α-helices on one side of the TOG domain comprise the microtubule interacting face (Ayaz et al., 2012; Byrnes & Slep, 2017). TOG domain-containing proteins usually have an array of multiple TOG domains with sequence divergence and distinct functions; TOG domains are classified into families based on sequence and structural similarities (Al-Bassam et al., 2007; Al-Bassam & Chang, 2011; Ayaz et al., 2012; Byrnes & Slep, 2017; Das et al., 2015; Leano & Slep, 2019). Broadly, TOG domains have preferential binding for either the ‘kinked’ α/β-tubulin heterodimer in the cytosol or at the ends of splayed protofilaments or the ‘straight’ α/β-tubulin heterodimers in the context of the microtubule lattice. Identifying the cellular or organismal repertoire of microtubule regulatory proteins is essential for understanding microtubule dynamics and regulation. The nematode *C. elegans*, with its simple body plan of diverse cell types and accessible genomics and genetics is a strategic model for this kind of study (Lacroix et al., 2014; Srayko et al., 2005).

To identify novel microtubule binding proteins in *C. elegans*, we searched its predicted proteome for TOG domain-containing proteins and identified two previously uncharacterized proteins. These proteins, which we named TOD-1 and TOD-2, are predicted by sequence similarity to be members of the XMAP215 family of microtubule polymerases (Al-Bassam et al., 2007; Brouhard et al., 2008; Byrnes & Slep, 2017; Widlund et al., 2011). These TOG domains are most similar to the TOG2 domains of the XMAP215 family (Al-Bassam & Chang, 2011). TOD-1 and TOD-2 are expressed within oogenic germline and sperm of adult hermaphrodites and in the germline of males. Animals from which each of these genes was deleted via genome editing were viable and generally healthy, although adult hermaphrodites laid unfertilized oocytes in an age-onset manner. This phenotype was due to abnormal sperm and not defects in the oogenic or somatic germline.

## Methods

### *C. elegans* strains, culturing, and mutagenesis

All *C. elegans* strains were maintained at 20°C using standard media and protocols (Brenner, 1974). All lines are described in Table S1.

CRISPR/Cas9 mediated mutagenesis was performed as described (Dokshin et al., 2018). A mix of crRNA, tracrRNA, Cas9, PRF4 (*rol-6(su1006)* extrachromosomal array), was injected into adult animals. Repair constructs were generated through mixing amplicons of mNeonGreen alone or mNeonGreen with long locus specific overhangs. To identify the optimal PAM site we used ‘Design CRISPR’ tool in the electronic lab notebook Benchling. gRNAs and oligonucleotides are listed in Table S2. Animals were confirmed by microscopy and were backcrossed to parental N2 wild-type animals three times.

### *C. elegans* live imaging conditions

Sample preparation: Animals were picked off plates and deposited in 5µL of M9 buffer containing 1mg/mL levamisole on a slide. Vaseline was deposited to serve as a spacer. A coverslip was gently pressed onto animals and slides were sealed with VaLaP.

Microscopes: Images were captured on two different systems. Images in Figure 2 were captured on a Nikon A1R microscope with a 60x 1.27 NA Nikon water immersion objective with a Gallium arsenide phosphide photo-multiplier tube (GaAsP PMT) detector using NIS-Elements. Image acquisition involved taking confocal sections with.5 µm spacing using the Galvano point scanner. Images of whole germlines required stitching of multiple frames using 15% overlap. Images in Figure 4 were captured on a Nikon Eclipse Ti2-FP widefield microscope with a Hamamatsu Orca FusionBT camera. Images were taken with either 20x, 0.75 NA air objective (Figure 4A,B) or with a 60x 1.20 NA water immersion objective (Figure 4C-E).

### Phylogenic analysis

TOG domain sequences from XMAP215, CLASP, or Crescerin family members were identified using the domain classification in UniProt. Protein sequence alignment was done in MEGA12 using MUSCLE and then a Maximum Likelihood Tree was constructed with a Jones-Taylor-Thornton model for substitutions (Jones et al., 1992; Saitou & Nei, 1987; Stecher et al., 2025).

### AlphaFold modeling

Sequences used for models were acquired from UniProt or WormBase. AlphaFold models were build using AlphaFold3. Models were opened in ChimeraX (Abramson et al., 2024; Meng et al., 2023; Pettersen et al., 2021). To thread TOG domains through the Stu2 structure on the kinked *S. cerevisiae* dimer, MatchMaker was used in default setting with the PDB file set as the template.

### Brood size assay

Animals in the final larval stage (L4) were placed onto individual plates and transferred every 24 hours to a fresh plate. This was done for 5 days. The number of hatched larva and embryos was counted 24 hours after the adult worm was moved to a fresh plate to ensure all embryos hatched. Total brood refers to the sum of the hatched larva and dead embryos. Embryonic viability was calculated by dividing the hatched larva by the total brood. The number of unfertilized oocytes was also counted. Example images of plates were taken using a Moticam X camera (Motic, California, USA) mounted on the eyepiece of a Nikon SMZ800 stereoscope microscope and recorded on an Apple iPhone 15 mobile phone. Experiment was repeated a total of three times for wild type, and two times for each mutant.

### Plate level phenotyping

Plates of gravid adults were washed with M9 and transferred to microcentrifuge tubes. Animals were pelleted and resuspended 1:1 in M9 and 2X bleach solution (2% bleach and.5 M NaOH). Tubes were vortexed until animals released embryos into solution. Animals were pelleted and bleach was removed and the pellet was resuspended in M9 and transferred to a new microcentrifuge tube and embryos were washed 3 times to removed residual bleach. Embryos were finally resuspended in M9 and rocked overnight at room temperature to hatch. Synchronized L1s were deposited onto growth plates with bacteria that allowed resumption of development. 3 days later the total number of animals was counted, and the visible phenotypes were scored. The occurrence of each phenotype incidence was calculated as occurrence divided by total animals counted. Experiment was repeated once with synchronized animals split across 3 plates for wild type and 5 plates for mutant animals.

### Oocyte cellularization assay

Animals in the final stage of larval development were placed on a fresh plate for 3 days (a timepoint where we see unfertilized oocytes laid based off the brood count). Animals were then mounted as described above and DIC images of both gonads were taken in a z-series. The number of cellularized compartments in the proximal germline was then counted. Experiment was repeated once.

### Sperm positioning assay

Otherwise wild-type or mutant animals with fluorescently labeled chromatin in the final larval stage were placed on fresh plates and at 24 hours or 48 hours were mounted and imaged. DIC and fluorescent images were taken of both gonad arms. Sperm nuclei were then counted and assigned as either in the proximal germline, spermatheca, or uterus. The fraction of sperm localized in the spermatheca, and proximal germline was then calculated by adding those two numbers together and then dividing by the total number of sperm. Experiment was repeated once with one biological replicate for wild type and two biological replicates for TOD-1 and TOD-2 knockouts.

### Mating assay

Female *fog-2(su1006)* L4 stage animals were incubated with excess wild-type or mutant male animals for 24 hours. Individual females were then placed singly onto growth plates and moved every 24 hours for 4 additional days. Like the brood size assay, hatched larvae and dead embryos were counted 24 hours after the adult was moved. Unfertilized oocytes laid on the plate were also counted. Total brood was the sum of hatched larvae and dead embryos; embryonic viability was calculated as hatched larva divided by the total brood. Experiment was repeated three times for each genotype.

### Figure preparation and statistics

All images were processed in FIJI before preparation in Adobe Illustrator (Schindelin et al., 2012). All graphs were made in GraphPad Prism. One-way ANOVAs were performed with a Tukey post-hoc test. For ANOVAs with a p < 0.05, a student’s t-test was performed to get an accurate p-value. Statistics were also done using a two-tailed analysis.

### AI Statement

All text written for this manuscript was human generated, no generative AI was used.

## Results

### Identification of uncharacterized TOG-domains in *C. elegans*

To identify novel regulators of microtubule dynamics in the model organism *Caenorhabditis elegans*, we searched for genes encoding common microtubule binding domains. We searched ‘TOG domain’ in InterPro to find all proteins classified as TOG domain-containing proteins (accession: IPR034085). InterPro defines TOG domains as part of a superfamily with Armadillo-like helical repeats; and have been identified due to sequence conservation (Andrade et al., 2001; Groves & Barford, 1999). We next limited our list of all TOG domain-containing proteins to those encoded by the *C. elegans* genome specifically. The resulting inventory included well-known and characterized genes: XMAP215^ZYG-9^, CLASP^CLS-1/CLS-2^, and Crescerin^CHE-12^ family members in the *C. elegans* genome (Figure 1A). We also identified two novel genes predicted to encode <u>TO</u>G <u>d</u>omains, T08D2.8 and F54A3.2, which we named TOD-1 and TOD-2. TOD-1 contains one TOG domain, unlike most TOG domain-containing proteins, and an S/R rich C-terminal region and is located on the X chromosome (Figure 1B). TOD-2 has two TOG domains and an S/R rich C-terminal region and is located on chromosome 2 (Figure 1B). To classify these novel TOG domains, we assessed the alignment of the TOG domains from XMAP215, CLASP, and Crescerin family members across five species, assembling a Maximum Likelihood Tree (Figure 1A; Jones et al., 1992; Saitou & Nei, 1987; Stecher et al., 2025). We also performed a BLAST search of TOD-1 and TOD-2 and only found gene family expansion within other *Caenorhabditis* species (data not shown). The TOG domains of TOD-1 and TOD-2 are most closely related to the TOG2 domains from XMAP215 family members ZYG-9, Msps, and CKAP5, in addition to the TOG1 domain from ZYG-9 (Figure 1A). XMAP215 family proteins are microtubule polymerases; our phylogeny suggests that TOD-1 and TOD-2 possess this function.

**Figure 1:**
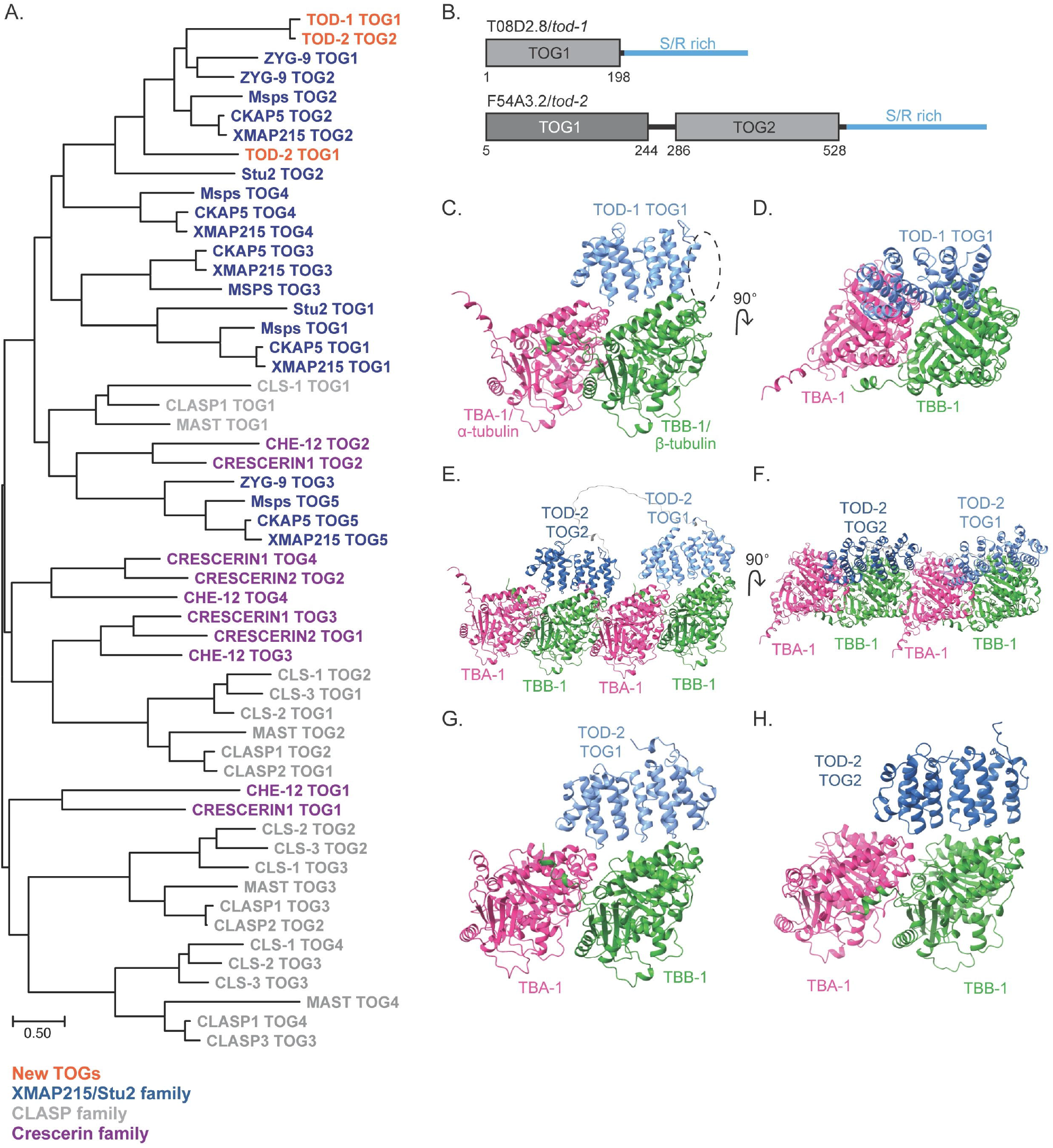
Identification of uncharacterized TOG domains in *C. elegans*. A. Maximum likelihood tree of XMAP215, CLASP, and Crescerin family members from: *C. elegans, Homo sapiens, Saccharomyces cerevisiae, Xenopus laevis*, and *Drosophila melanogaster*. TOG domains are color-coded by family: XMAP215 in blue, CLASP in grey, Crescerin in purple and novel TOG domains in orange. B. Domain structure for T08D2.8/TOD-1 and F54A3.2/TOD-2. TOG1 of TOD-1 and TOG2 of TOD-2 are in light gray, TOG1 of TOD-2 is in light gray, and Serine/Arginine-Rich region in blue. C. AlphaFold model of TOD-1 on a TBA-1/TBB-1 dimer. Looking along the protofilament. Missing heat repeat is outlined with black circle. D. 90° turn of AlphaFold model from panel C. Looking down onto the surface of a microtubule. E. AlphaFold model of TOG1 and TOG2 domains of TOD-2 on two TBA-1/TBB-1 dimers. View looking along protofilament. F. 90° turn of alpha fold model from panel E. Looking down onto the surface of a microtubule. G, H. AlphaFold model of TOG1 (G) and TOG2 (H) domains of F54A3.2/TOD-2 on a TBA-1/TBB-1 dimer. Side view of protofilament.

We next used AlphaFold3 to predict how each protein interacts with a dimer of *C. elegans* tubulin (Abramson et al., 2024). The single TOG1 domain from *tod-1* is predicted to interact with the tubulin dimer similarly to how other XMAP215 family members do, through conserved amino acids in the intra-HEAT loops positioned along one face of the α-solenoid structure (Figure 1C, D; Al-Bassam et al., 2007; Byrnes & Slep, 2017). Interestingly, the single TOG domain in TOD-1 only has five HEAT repeats and not the typical six. The missing HEAT repeat would bind towards the plus end of the microtubule (Figure 1C, noted in dashed, black oval). The TOG domains in TOD-2, like TOD-1, are both predicted to interact with tubulin dimers as other XMAP215 family proteins (Figure 1E-H). One key difference is that XMAP215 family members with more than two TOG domains also contain TOG domains with a lattice binding preference (Al-Bassam & Chang, 2011). To identify if the novel TOG domains have a similar domain structure to the TOG2 domain of Stu2, we used the Matchmaker function in ChimeraX to thread each TOD protein TOG domain sequence through the TOG2 domain of the *Saccharomyces cerevisiae* protein Stu2 on the ‘kinked’ (soluble) conformation of a tubulin dimer (Figure S1; Meng et al., 2023; Pettersen et al., 2021). The TOG domains of TOD-1 and TOD-2 aligned to the kinked conformation of tubulin (Figure S1). Together these data suggest that the single TOG domain of TOD-1 and both TOG domains of TOD-2 bind not to the microtubule lattice, but to soluble tubulin.

### *tod-1* and *tod-2* are expressed in the sperm

To determine where TOD-1 and TOD-2 are expressed and localize, we generated fluorescently tagged lines using CRISPR/Cas9 mediated genome editing (Dokshin et al., 2018). mNeonGreen was added to the C-terminal end of both *tod-1* and *tod-2*. Animals were then imaged 24 hours after the final larval stage. We found, consistent with mRNA sequencing data, that both TOD-1 and TOD-2 are expressed at low levels in a variety of tissues (Ghanta et al., 2021; Packer et al., 2019). TOD-1 was most notably present in the sperm (Figure 2A). TOD-2-mNG intensity was stronger among different tissues and cell types compared to TOD-1 (Figure 2B). TOD-2 was most abundant in the sperm of adult hermaphrodites and males (Figure 2B, C). Curiously, not all sperm appeared to have the same abundance of the novel proteins. This expression pattern suggests that deletion of either *tod-1* or *tod-2* may affect fertility.

**Figure 2:**
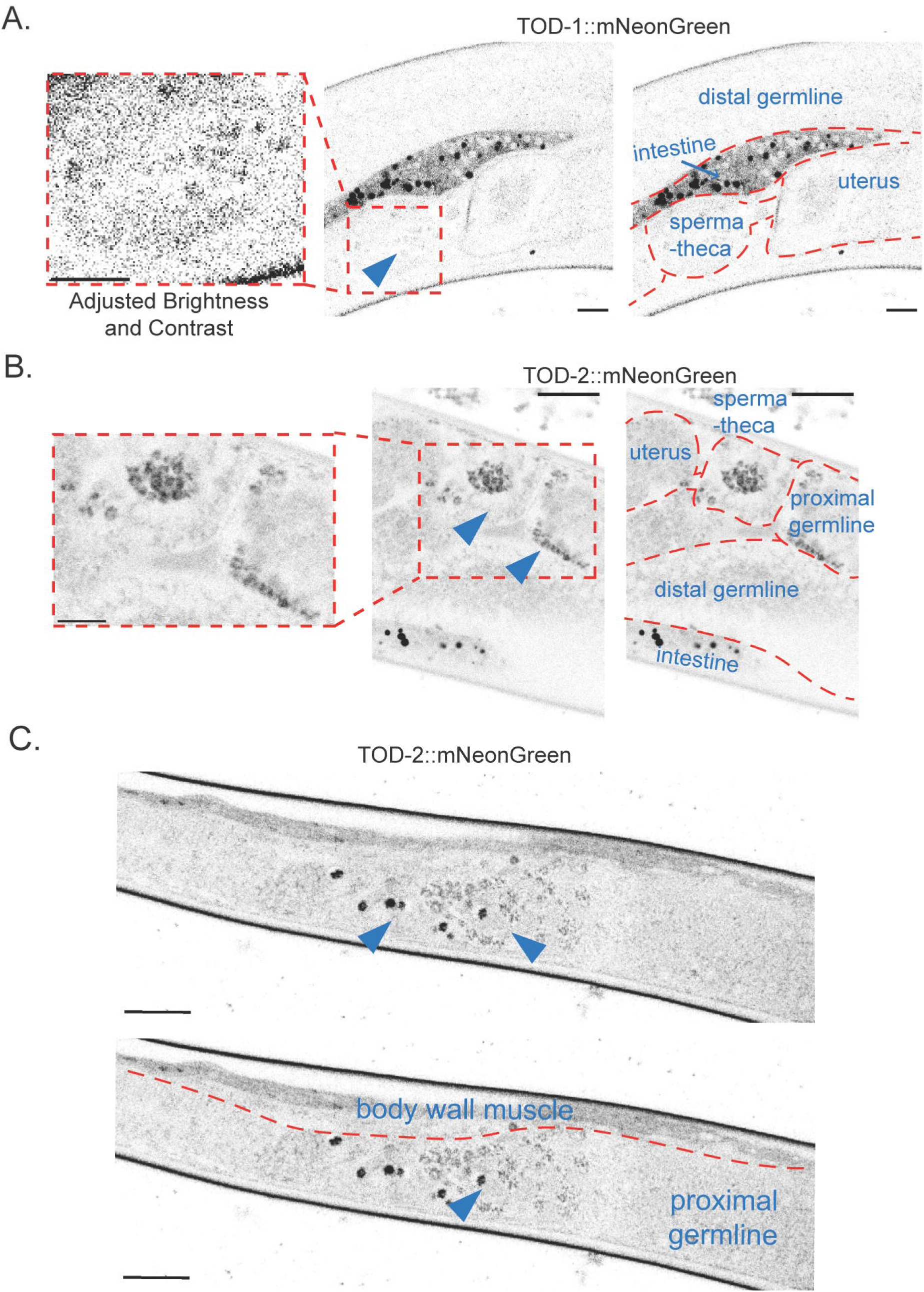
TOD-1 and TOD-2 are enriched in sperm. A. TOD-1::mNeonGreen in an adult hermaphrodite. Zoom in on the far left with adjusted brightness and contrast. Outline on the right to indicate organs. Scale bar equals 10 microns. B. TOD-2::mNeonGreen in an adult hermaphrodite. Zoom in on the far left. Outline on the right to indicate organs. Scale bar: 10 microns. C. TOD-2::mNeonGreen expression in an adult male. Zoom in on the far left. Outline on the right to indicate organs. Scale bar: 10 microns.

### *tod-1* and *tod-2* are dispensable for fertility

We next tested how knocking out *tod-1* and *tod-2* impacted animal fertility and physiology. To do this we replaced the open reading frame of *tod-1* and *tod-2* with mNeonGreen, creating both a transcriptional reporter and a knockout. We next scored phenotypes visible on a stereomicroscope, synchronizing animals during the first larval stage and examining them after 3 days. We then counted the total number of animals on each plate and scored animals for visible phenotypes. Two phenotypes occurred with low incidence: immature animals that were still in the last larval stage, and vulval protrusion (Pvl), indicating perturbation of tissue biogenesis or maintenance of tissue integrity (Fay, 2013; Perry et al., 2023) (Figure 3A). We found that approximately 10% of *tod-1* and *tod-2* knockouts were still in the final stage of larval development compared to 2% of the wild-type animals (Figure 3A). 7.5% of TOD-2 knockouts were Pvl (Figure 3A). Stereoscope-accessible phenotypes that were not observed included dumpy (short) or long body, cuticle blistering, and uncoordinated motility. All together, these data demonstrate that TOD-1 and TOD-2 are only stochastically and rarely required for *C. elegans* postembryonic development, but knockouts do increase the occurrence of unfertilized oocytes being laid.

**Figure 3:**
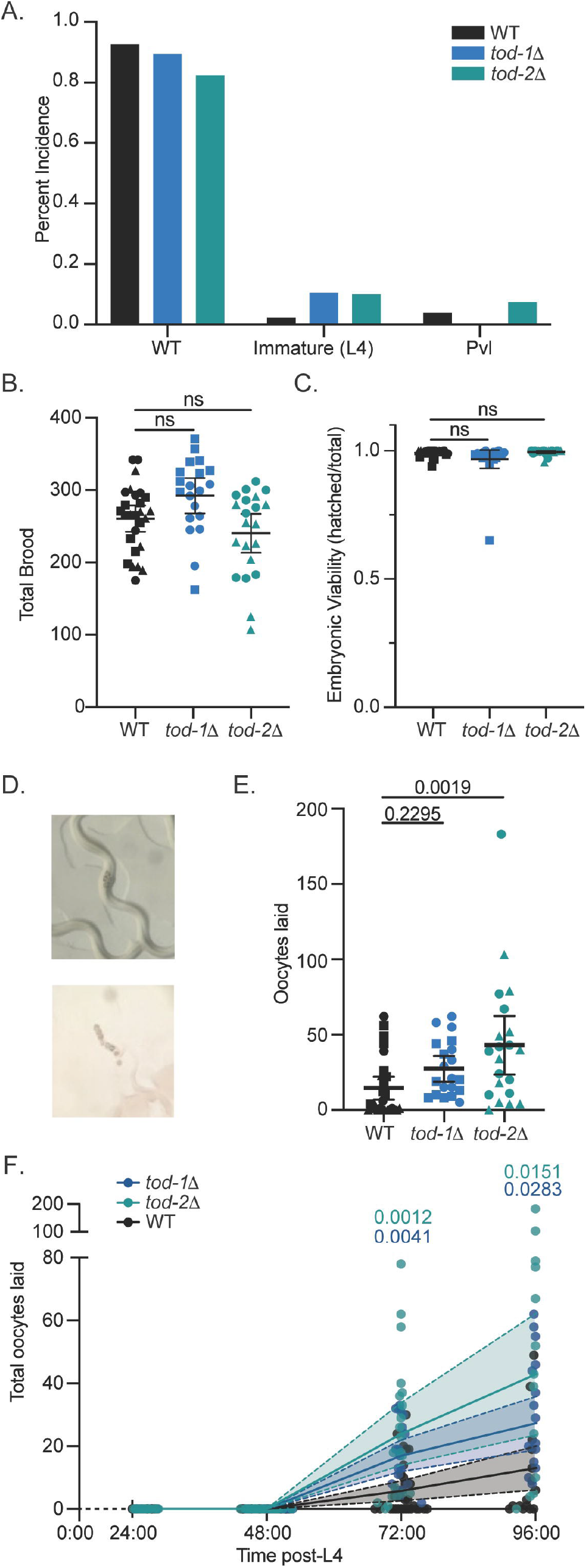
TOD-1 and TOD-2 are dispensable for fertility but necessary to avoid laying unfertilized eggs. A. Plate level phenotyping for wild type (n=176 animals), *tod-1*Δ (n=123 animals), and *tod-2*Δ (n=200 animals). Plates were scored for immature animals and animals with protruded vulva (Pvl). B. Total brood count for wild type (n=27), *tod-1*Δ (n=21), and *tod-2*Δ (n=20). Total brood was measured over 5 days. Bar is mean error bars are 95% confidence interval. Adjusted P-values were calculated from the post hoc test after a one-way ANOVA. Each dot represents one hermaphrodite and shape represents experimental replicate. C. Embryonic viability of wild type (n= 27), *tod-1*Δ (n=21), and *tod-2*Δ (n=20). Ratio was calculated as total hatched animals/total brood from blood count. Bar is the mean; error bars are 95% confidence interval each dot represents one hermaphrodite and shape represents experimental replicate. P-values were calculated from post hoc tests after one way ANOVA. D. Example images of unfertilized oocytes on plates. E. Total number of unfertilized oocytes during 5-day blood count of wild type (n=27), *tod-1*Δ (n=21), and *tod-2*Δ (n=20). Bar is the mean; error bars are 95% confidence interval. Each dot represents 1 hermaphrodite and shape represents experimental replicate. True P-values are from a student’s T test after a one-way ANOVA. F. Cumulative number of unfertilized oocytes laid during 4-day blood count of wild type (n=27), *tod-1*Δ (n=21), and *tod-2*Δ (n=20) animals. Dot: individual animal. Adjusted P-values are from a post hoc test after a two-way ANOVA.

We next assessed how knocking out each of these genes impacts fertility by counting hermaphrodites’ total brood. We found no change in the total brood count (sum of hatched animals and unhatched embryos) when either *tod-1* or *tod-2* is knocked out compared to wild-type control animals (Figure 3B). *tod-1* knockouts had a slight increase in total brood, while *tod-2* knockouts had a slightly smaller brood count compared to wild-type animals, but these differences were not statistically significantly different (Figure 3B).

We next asked if *tod-1* or *tod-2* knockout affects embryonic viability. There was no change in the fraction of embryos that hatch in *tod-1* or *tod-2* knockout animals versus controls (Figure 3C). Inspection of singled knockout animals’ growth plates revealed that these animals laid unfertilized oocytes (Figure 3D, E). Unfertilized oocytes are circular and lack the eggshell and thus optical contrast of an embryo and thus appear dark on the plate (Figure 3D). Counting these oocytes revealed that knockout of either *tod-1* or *tod-2* resulted in an increased number of unfertilized oocytes laid by adult hermaphrodites (Figure 3D, E). Laying of unfertilized oocytes also results from mutation in sperm proteins SPE-6 *and* SPE-38; such abnormal reproductive output occurs steadily throughout adulthood in these conditions (Chatterjee et al., 2005; L’Hernault et al., 1988a; Peterson et al., 2021). We examined the timing of laying unfertilized oocytes in *tod-1* and *tod-2* knockouts, and determined that they exhibit this phenotype in an age onset manner (Figure 3F). Unlike the spe mutants, *tod-1* and *tod-2* null worms lay unfertilized oocytes in an age-onset manner (Figure 3F). Both *tod-1* and *tod-2* knockouts lay more unfertilized oocytes at early timepoints compared to wild-type animals (Figure 3F). These data suggest that TOD-1 and TOD-2 are dispensable for fertility but required for retention of unfertilized oocytes.

### *tod-1* or *tod-2* deletion impairs normal sperm physiology

*C. elegans* hermaphrodites generate and fertilize oocytes, and lay embryos, as long as sperm are present in the spermatheca. After the sperm supply has been exhausted, oogenesis continues but cellularized oocytes “stack up” in the proximal germline. We hypothesized that *tod-1* and *tod-2* knockout animals lay unfertilized oocytes due to defects in the crosstalk between sperm and the oogenic germline. To test this idea, we captured DIC images of wild-type or mutant animals at a timepoint when unfertilized oocytes were laid: 48 hours after the final larval stage. We counted the number of cellularized compartments proximal to the spermatheca (Figure 4A, red bracket). *tod-1* and *tod-2* knockouts had 1.24- and 1.29-fold more cellularized oocytes in the distal germline than wild type, respectively (Figure 4B). Interestingly, the presence of unfertilized oocytes was often different between the anterior and posterior gonads (Figure 4A). These data suggest fertilization is impaired in *tod-1* and *tod-2* mutant animals.

**Figure 4:**
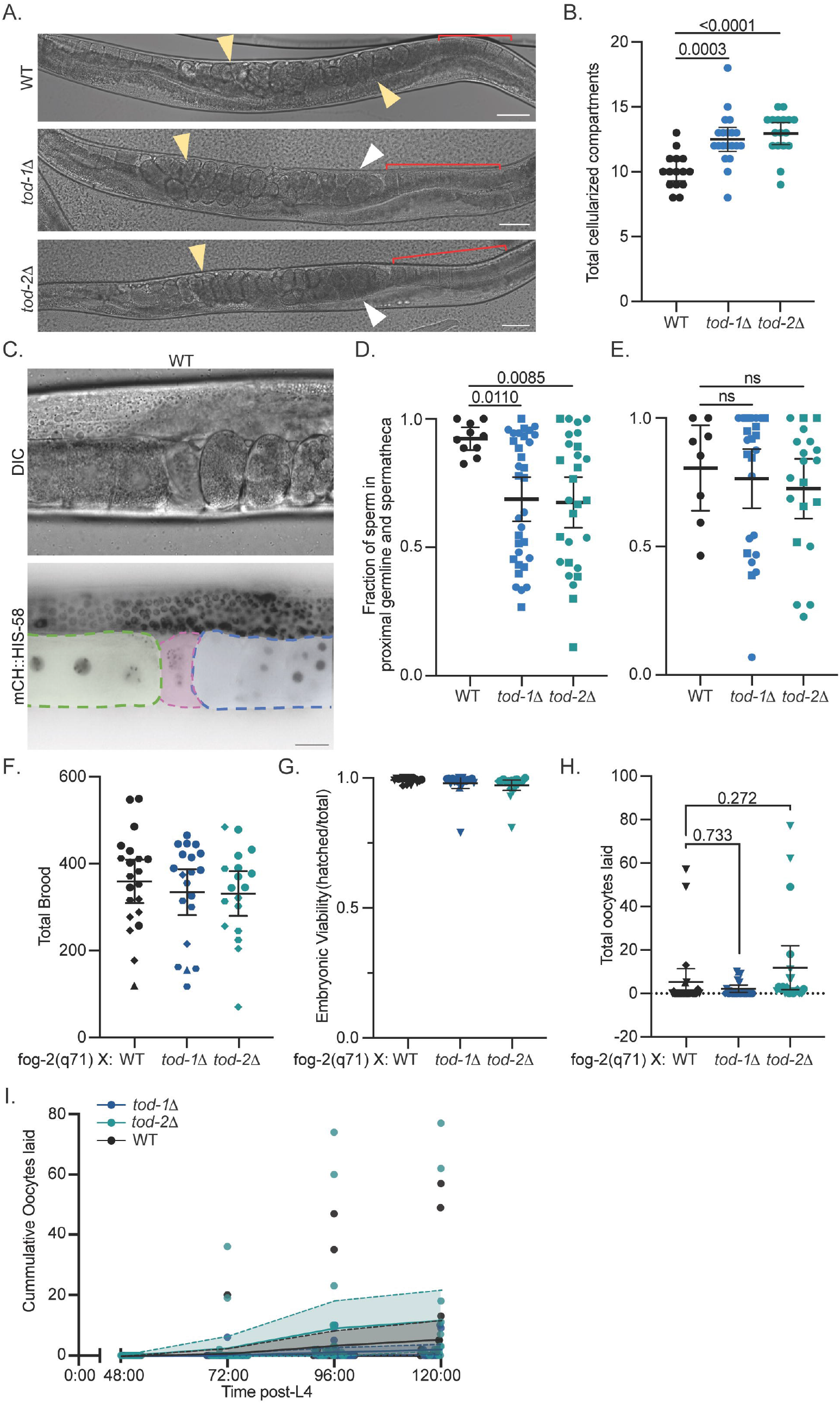
TOD-1 and TOD-2 promote sperm function. A. Example DIC images of wild type, *tod-1*Δ, or *tod-2*Δ animals. Yellow arrowheads: embryos; white arrowheads: unfertilized oocytes. Red bracket: cellularized compartments quantified. B. Total number of cellularized compartments from wild type (n=15), *tod-1*Δ (n=17), or *tod-2*Δ (n=20) animals. Bar: mean; error bars: 95% confidence intervals. P-values are from a student’s T-test after a one-way ANOVA. C. Example DIC and fluorescence image of a wild-type animal. Green area outlines proximal germ line, pink area web outlines spermatheca, and blue area outlines the uterus. Scale bar is 5µm. D. Fraction of sperm localized to the spermatheca and proximal germ line 24 hours after the final larval stage for wild type (n=10), *tod-1*Δ (n=33), or *tod-2*Δ (n=28) animals. Bar: mean; error bars: 95% confidence intervals. Each dot represents one animal and shape represents biological replicate. P-values are from a student’s T-test after a one-way ANOVA. E. Fraction of sperm localized to the spermatheca and proximal germline 48 hours after the final larval stage(n=8), *tod-1*Δ (n=20), or *tod-2*Δ (n=24) animals. Bars is the mean; error bars are 95% confidence interval. Each dot represents one animal shape represents biological replicate. Adjusted P-values are from a post hoc test after a one-way ANOVA. F. Total brood count for *fog-2(q71)* female animals mated with wild type (n=21), *tod-1*Δ (n=20), or *tod-2*Δ (n=18) males. Brood count was measured from 24 hours after the final larval stage to 5 days after the final larval stage. Bars is the mean; error bars are 95% confidence interval. Each dot represents one animal and shape represents experimental replicate. Adjusted P-values are from a post hoc test after a one-way ANOVA. G. Embryonic viability for *fog-2(q71)* female animals mated with wild type (n=21), *tod-1*Δ (n=20), or *tod-2*Δ (n=18). Ratio was calculated as total hatched animals/total brood from blood count. Bar is the mean; error bars are 95% confidence interval each dot represents one hermaphrodite and shape represents experimental replicate. P-values were calculated from post hoc tests after one way ANOVA. H. Total number of unfertilized oocytes laid during 4-day blood count of *fog-2(q71)* female animals mated with wild-type (n=21), *tod-1*Δ (n=20), or *tod-2*Δ (n=18) males. Bar is the mean; error bars are 95% confidence interval. Each dot represents 1 hermaphrodite. Adjusted P-values are from a student’s T test after a one-way ANOVA. I. Cumulative number of unfertilized oocytes laid during 4-day blood count of *fog-2(q71)* female animals mated with wild-type (n=21), *tod-1*Δ (n=20), or *tod-2*Δ (n=18) males. Dot: individual animal.

During fertilization in *C. elegans*, as the embryo moves from the spermatheca into the uterus, sperm are also displaced into the uterus. Because sperm and the oocyte must be together in the spermatheca for fertilization to take place, displaced sperm must crawl back into the spermatheca. Defects in sperm motility and positioning can result in reduced brood count and the laying of unfertilized oocytes (Geldziler et al., 2011; Mei & Singson, 2021; Singson, 2001). We hypothesized that the increased number of unfertilized oocytes laid is due to a decreased availability of sperm capable of fertilizing oocytes, due to, for example, inability of sperm to return from the uterus to the spermatheca (Figure 4C, blue outline). To assess this, we labeled the nuclei of wild-type and mutant animals with an mCherry tag on histone H2B and imaged adult animals 24 and 48 hours after the final larval stage. We then quantified the number of sperm localized in the spermatheca-proximal arm of the oogenic germline (Figure 4C, green outline), in the spermatheca (pink outline), and in the uterus (blue outline) and calculated the fraction of sperm localized within the proximal germline and spermatheca. At 24 hours after the final larval stage, both *tod-1* and *tod-2* knockout animals had a lower fraction of sperm localized in the proximal germline and spermatheca than controls (Figure 4D). Interestingly, at 48 hours after the final larval stage, *tod-1* and *tod-2* knockout animals’ sperm mobility/homing was similar to wild type (Figure 4E). This result suggests that TOD-1 and TOD-2 are both required for sperm homing to the spermatheca.

Return of sperm to the spermatheca from the uterus not only requires sperm motility but is also dependent on the release of homing signals by oocytes (Kubagawa et al., 2006). Therefore, we tested if the increased laying of unfertilized oocytes by *tod-1* and *tod-2* knockout animals was due to a defect in sperm or whether TOD-1 and/or TOD-2 are also required in the somatic gonad or oocytes. To do so, we used a *C. elegans* mutant, *fog-2(q71)*, that only produces male and female animals. Wild-type or *tod-1* or *tod-2* knockout males with labeled nuclei were mated in excess to the *fog-2(q71)* females in the final larval stage for 24 hours. We then assessed the fecundity of the mated *fog-2(q71)* adult females. As with animals in which all somatic and germline tissues bear the TOD-1 and TOD-2 knockout, the total brood count for *fog-2(q71)* females mated with TOD-1 and TOD-2 knockout males was indistinguishable from those mated with wild-type males (Figure 4F). Further, embryos fertilized with either *tod-1* or *tod-2* knockout sperm had normal viability (Figure 4G). However, *fog-2(q71)* females mated with *tod-1* or *tod-2* knockout males laid a few more unfertilized oocytes than those mated with wild-type males (Figure 4H). Like our full knockout animals, *fog-2(q71)* females mated to *tod-1* or *tod-2* males laid unfertilized oocytes in an age onset manner (Figure 4I). However, the *fog-2(q71)* animals mated with *tod-1* or *tod-2* knockout sperm exhibited a slight but not statistically significantly early onset laying of oocytes compared to wild-type animals (Figure 4I). Together, our findings suggest that TOD-1 and TOD-2 play a role in normal sperm function but that the unfertilized oocyte phenotype is not due solely to each protein’s function in sperm.

## Discussion

We identified two previously uncharacterized TOG domain-containing proteins. Sequence and structural conservation classified these proteins in the XMAP215 family of TOG domain-containing proteins and predicted they bind to the soluble tubulin heterodimers. Microtubule lattice-binding by the S/R rich tail of each TOD protein may allow the dimer-binding TOG domain(s) to promote microtubule polarization (Figure 1A, B S1; (Al-Bassam & Chang, 2011; Currie et al., 2011; Geyer et al., 2018; Widlund et al., 2011) Future work will test whether TOD-2 binds microtubules and concentrates dimers there to promote polymerization like a canonical XMAP215 family member.

TOD-1 is unique among TOG domain-containing proteins for two reasons. First, it contains only one TOG domain, making it the first such animal TOG domain-containing protein; the only other known single TOG domain-containing protein is SPIRAL2 in *Arabidopsis thaliana* (Fan et al., 2024). A defining feature of SPIRAL2 is that it is a hexamer in the cytosol and recruits katanin to microtubules to drive severing (Fan et al., 2024). TOD-1 only contains a TOG1 domain and an S/R-rich C-terminal tail that is not predicted to oligomerize (data not shown). Furthermore, the TOG1 domain of TOD-1 only contains five HEAT repeats, not the typical six (Figure 1). The “missing” HEAT repeat would interact towards the exposed “plus-end” of the β-tubulin of a dimer (Figure 1). Future work characterizing the activity of this single, truncated TOG domain protein with no predicted oligomerization domain will expand the “rules” of how XMAP215 family members functions.

TOD-1 and TOD-2 are expressed in multiple tissues and are most abundant in the sperm of adult animals (Figure 2). Both *tod-1* and *tod-2* are expressed at low (mRNA) levels; indeed the proteins are of low abundance (Figure 2; Ghaddar et al., 2023; Packer et al., 2019). The low abundance of TOD-1 and TOD-2 in large cells such as the uterine muscles or early blastomeres and relative enrichment in sperm that are thought to lack microtubules precluded our ability to visualize them on individual microtubules in *C. elegans* (Hu et al., 2019; Shakes et al., 2009). By contrast, ZYG-9 is visible along meiotic and mitotic spindles (Bellanger et al., 2007; Bellanger & Gönczy, 2003; Cavin-Meza et al., 2022; Matthews et al., 1998). Identifying how TOD-1 and TOD-2 interact with α/β-tubulin heterodimers and microtubules and modulate microtubule dynamics will require further investigation.

Animal physiology was largely normal when either *tod-1* or *tod-2* was deleted (Figure 3). Mutant animals laid an abnormally high number of unfertilized oocytes compared to wild-type animals. This behavior is predicted to reduce animal fitness, since oogenesis is metabolically costly (L’Hernault et al., 1988a; Minniti et al., 1996; Peterson et al., 2021). Other mutants that produce an increased number of unfertilized oocytes have reduced total brood count, unlike our results (L’Hernault et al., 1988b; Mei & Singson, 2021; Minniti et al., 1996; Singson, 2001). We identified that the unfertilized oocyte laying phenotype is partially due to a defect in sperm and not due to defects in the oogenic germline or somatic gonad (Figure 4C-H). *C. elegans* sperm motility is independent of the microtubule or actin cytoskeletons, instead utilizing major sperm protein. There is no known role for, nor evidence of, microtubules in *C. elegans* sperm; it is thought that microtubules are eliminated from sperm, relegated to the residual body during spermatogenesis (Shakes et al., 2009; Shakes & Ward, 1989). We speculate that TOG domain-containing proteins impact sperm activity by inhibiting the polymerization of any residual tubulin, or limiting the length of microtubules in sperm that are functionally important for some aspect of cell biology such as scaffolding organelles, indirectly impacting fertilization or motility. We previously identified homologs of the heterodimer binding proteins cylicin, CYLC-1 and CYLC-2, as highly enriched in sperm (Krauchunas et al., 2020; Lacroix et al., 2014, 2016). Future work will be aimed at uncovering the roles of these two different families of tubulin heterodimer binding proteins in *C. elegans* sperm.

Future work will be aimed at determining how these novel XMAP215 family proteins impact microtubule dynamics. How does TOG domain number change microtubule dynamics? What defines a TOG domain’s ability to promote polymerization? Are there genetic or physical interactions among XMAP215 family members to promote or oppose polymerase activity? How does the single, divergent TOG domain of TOD-1 regulate microtubules? The novel proteins we report here provide opportunities to explore and identify the defining characteristics and rules for TOG domain-containing proteins’ structural and functional diversity.

## Acknowledgements

This work was supported by NSF 2153790 to ASM. LCW was supported by the NIH grant 5K12GM000678 (PIs: Donald T Lysle and Kathryn J Reissner). We are grateful for Kacy Gordon’s lab for providing the *fog-2(q71)* line. We would like to thank Kevin Slep and Udo Onwubiko for careful reading and comment on the manuscript. We would also like to thank members of the Maddox labs for helpful discussion about this project, particularly Michael Werner.

## Figure Legends

**Figure S1:**
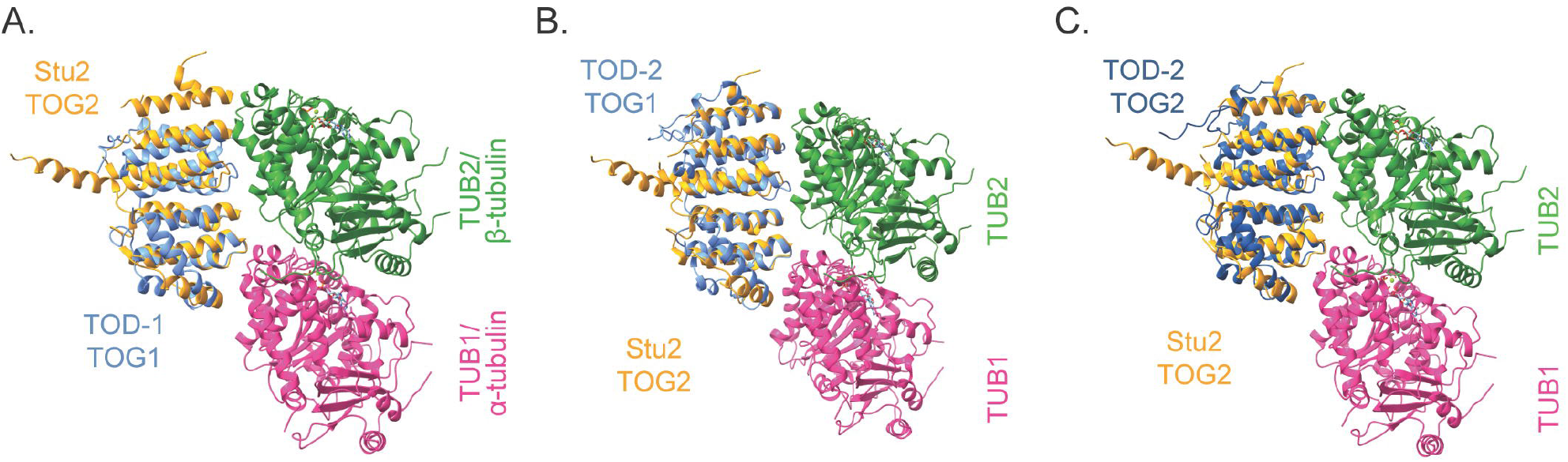
*tod-1* and *tod-2* align to the Stu2 TOG2 domain. A. TOG1 domain of *tod-1* aligned to Stu2 TOG2 domain from *S. cerevisiae* using Matchmaker in ChimeraX. Structure from PDB: 4FFB. B. TOG1 domain of *tod-2* aligned to Stu2 TOG2 domain from *S. cerevisiae* using Matchmaker in ChimeraX. Structure from PDB: 4FFB. C. TOG2 domain of *tod-2* aligned to Stu2 TOG2 domain from *S. cerevisiae* using Matchmaker in ChimeraX. Structure from PDB: 4FFB.

### Strain tabl

| Strain name | Genotype | Source |
| --- | --- | --- |
| CB4108 | fog-2(q71) V. | Gift from Kacy Gordon's lab |
| MDX47 | ItIs37 [pAA64; pie-1/mCHERRY::his-58; unc-119 (+)] IV | (Smith, 2018) |
| MDX179 | mon31 (F54A3.2::mNG) II. | This study |
| MDX190 | mon32 (F54A3.2Δ::mNG) II. | This study |
| MDX193 | mon33 (T08D2.8Δ::mNG) X. | This study |
| MDX199 | ItIs37 [pAA64; pie-1/mCHERRY::his-58; unc-119 (+)] IV; mon32 (F54A3.2Δ::mNG) II. | This study |
| MDX201 | ItIs37 [pAA64; pie-1/mCHERRY::his-58; unc-119 (+)] IV; mon33 (T08D2.8Δ::mNG) X. | This study |
| MDX203 | mon36 T08D2.8::mNG X. | This study |

### Oligonucleotides and guideRNAs

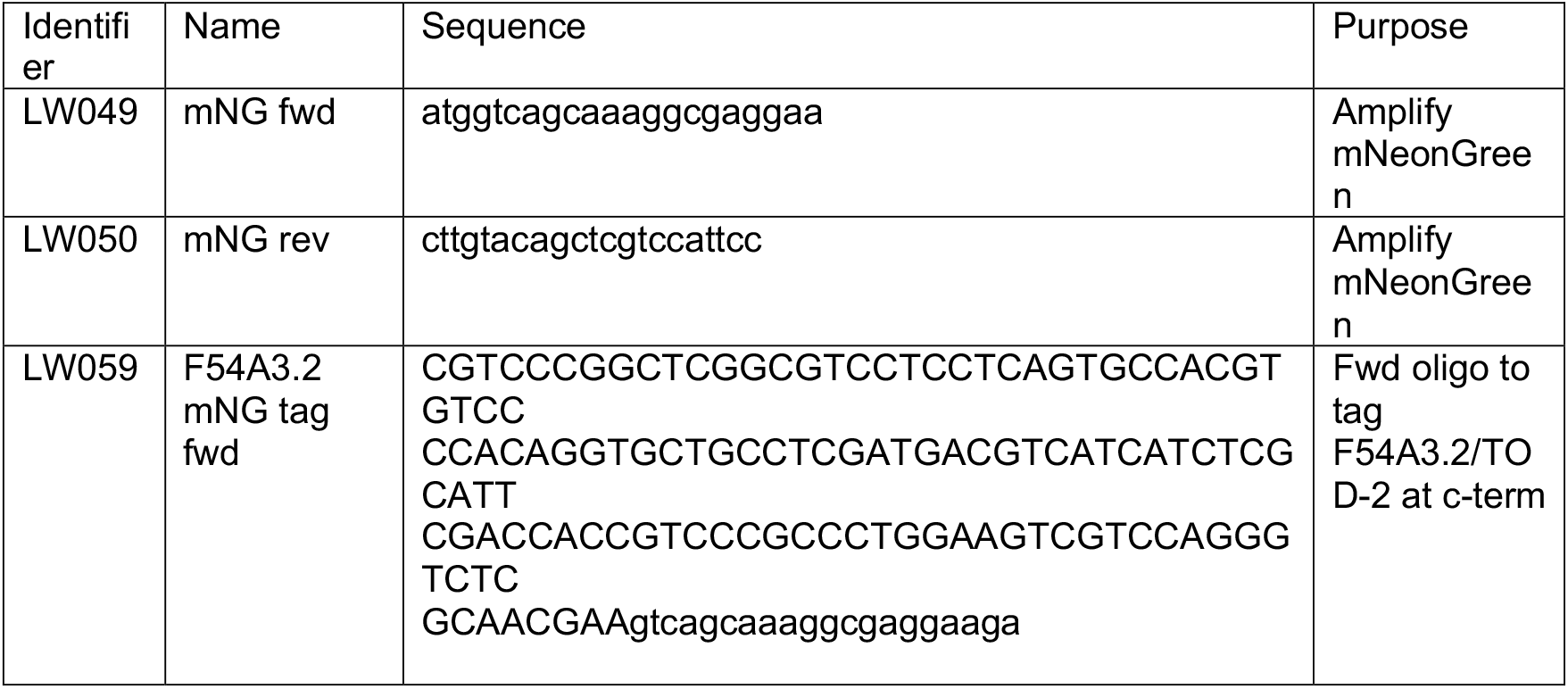
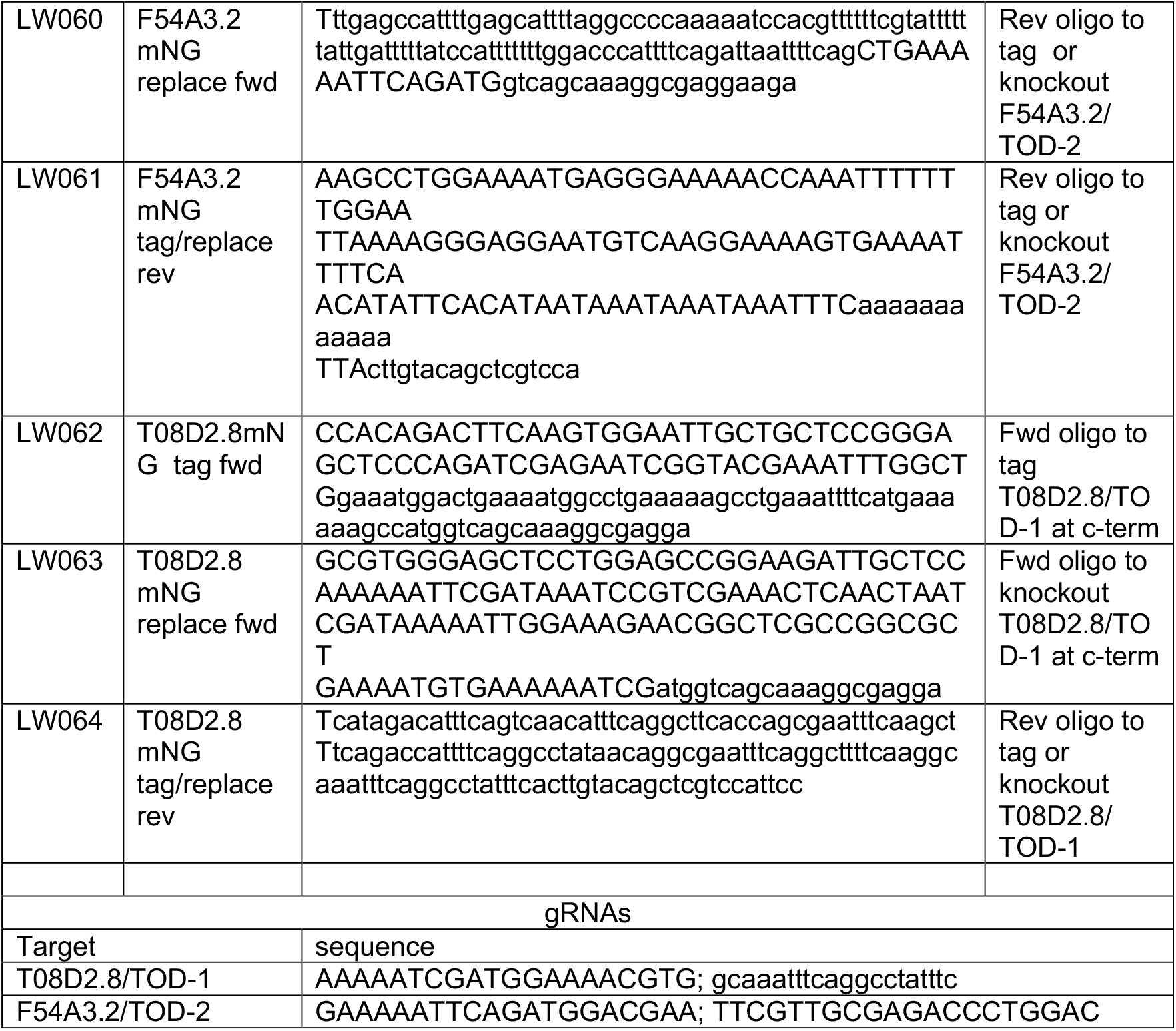

